# How the liver contributes to stomach warming in the endothermic white shark *Carcharodon carcharias*

**DOI:** 10.1101/599639

**Authors:** David C. Bernvi, Geremy Cliff

**Author notes:** Correspondence: David C. Bernvi, Hornsbergs Strand 21, 112 17, Stockholm, Sweden, e-mail: (DCB), (GC).

## Abstract

**Background:** White sharks and other lamnids are able to elevate their stomach temperature. The ability to heat large amounts of food to the recorded levels of up to 17°C above the ambient water temperature can’t be explained only by the heat generated by catabolism and the endothermic adaptions of the suprahepatic rete. This rete has two shunts that regulates the volume of blood flowing from the liver to the sinus venosus, thereby providing a temperature control mechanism for the GIT. The liver’s function in this temperature regulation is unknown. White shark stomach volume is well above 135 l in sub-adults to accommodate large prey items, including marine mammals. The simultaneous ingestion of large volumes of cold water during feeding will result in marked cooling of the stomach. Our study investigated the role of the liver in relation to warming the gastro-intestinal tract and the retention of elevated core temperatures.

**Materials and methods:** The liver morphology and its position relative to the gastro-intestinal tract were investigated in 13 white sharks *Carcharodon carcharias*. Stomach volume and the thickness of the abdominal wall were also measured to give a comparative estimate of heat insulation by white muscles.

**Results:** In all white sharks examined the two liver lobes completely enveloped the entire gastrointestinal tract, with the anterio-ventral margins of the liver almost interlocking around the stomach. A large, conspicuous, flattened vascular system was only present on the inner surfaces of both liver lobes. The thickness of the ventral abdominal body wall is only 12% of that of the dorso-lateral body wall, so the potential for heat loss from the GIT via the belly region is high.

**Conclusion:** Our study builds on the findings of other researchers which revealed that the liver and digestive tract receive a major portion of their blood supply through the suprahepatic rete, which is a heat exchanger aimed at retaining metabolic heat generated by the red locomotory muscles. This heat is not only transferred to the stomach via its supply of warm blood but also via thermal conduction from the vessels on the inside of the liver, which envelopes the digestive tract and serves as a large reservoir of venous blood. The liver is rich in lipids, with insulatory properties to retain the heat which would otherwise be rapidly lost through the extremely thin ventral abdominal wall in temperate waters, where white sharks commonly occur. These findings provide insight into the hitherto unknown role played by the liver in the highly elevated stomach temperatures reported, thereby providing this endothermic top predator with enhanced rates of digestion.

## Introduction

In lamnid sharks (family Lamnidae) and the common thresher shark *Alopias vulpinus*, conservation of body heat, principally generated by red muscles (Carey & Teal 1969, Sepulveda *et al*. 2005), is facilitated by the location of the *retia mirabilia* between the red muscles and the subcutaneous lateral vessels (Carey *et al*. 1985). Cold, oxygenated blood from the gills enters the red muscles where it is warmed by deoxygenated blood flowing from the body core to the heart (Carey *et al*. 1981).

An elevated body temperature increases metabolic activity (Carey 1982); this is highly beneficial for species such as the white shark *Carcharodon carcharias* which inhabits cooler temperate waters. The benefits include increased power output from the muscles, shorter recovery time after bursts of speed, increased aerobic metabolic rate leading to faster digestion and internal temperature regulation (Carey 1982, Goldman 1997). Temperatures within the epaxial musculature may reach 3-5°C above the ambient water (absolute internal temperatures of 18-21°C) for individuals of 3.5-4.6 m (Carey *et al*. 1982, Tricas & McCosker 1984). White sharks of similar sizes also have an absolute elevated stomach temperature of about 26°C (Goldman 1997, Goldman *et al*. 1996, Lowe & Goldman 2001, McCosker 1987), reaching up to 17°C above the ambient water (Hoyos-Padilla *et al*. 2016), thereby greatly accelerating digestion of food.

The red muscles are known to generate heat, however it is believed that the elevated visceral temperatures observed in lamnids are generated from catabolism via digestion rather than heat generated by the red muscles and that the suprahepatic rete aids to conserve this catabolic heat (Bernal *et al*. 2011). A venous passage leading through the suprahepatic rete directly from the liver to the sinus venosus, thereby bypassing this visceral rete, was documented by Burne (1923), suggesting there is a mechanism for heat regulation (Carey *et al*. 1981). The hepatic sinus formed in the liver is connected by hepatic veins to the suprahepatic rete (Bernal *et al*. 2011) and may be closed by sphincter muscles, forming two shunts of the blood flow (Carey *et al*. 1981). Similar sphincter muscles in the hepatic veins regulate the blood flow from the liver to the heart in elasmobranchs (Johansen & Hanson 1967). Hence the hepatic sinus shunts regulates the temperature of the GIT in lamnids by allowing warm venous blood from the liver to flow through the suprahepatic rete rather than directly to the heart (Carey *et al*. 1981) and thereby warming the GIT.

When the hepatic sinus shunts are closed, it is possible that the liver may function as an organ for heat distribution. The contraction of the shunts would result in an increased amount of venous blood in the liver. Hence the heat retention in nearby organs such as the stomach would increase. Bernal *et al*. (2011) state that the potential role played by organs such the liver in the endothermy of white sharks and other lamnids has not been investigated and remains unknown. The objective of this study was to investigate the position and anatomy of the liver in relation to the stomach and intestine. The study also provided an opportunity to measure the abdominal body wall thickness which is known to vary among the endothermic lamnids and to constitute a potential site of heat loss (Carey *et al*. 1985).

## Materials and methods

### Specimens examined

All 13 white sharks examined in this study were caught in beach protection nets and drumlines maintained on the east coast of South Africa by the KwaZulu-Natal Sharks Board (KZNSB) (Cliff *et al*. 1989, Cliff and Dudley 1992, Cliff & Dudley 1996, Cliff & Dudley 2011). The sharks were frozen prior to examination. Based on the size classes of Hussey *et al*. (2012), they were all classified as juveniles except the largest, a sub-adult.

### Liver

The bi-lobed liver was photographed *in situ*, removed and weighed. To show the vascularisation on the inside of the liver lobes, liquid latex was injected into the two largest vessels connected to the suprahepatic rete of one of the sharks (RB15017).

### Stomach volume

The stomach was removed with the oesophagus and duodenum attached. Any contents were removed by temporarily everting the stomach and washing it with a fine spray of water. The pyloric canal was closed using a cable tie. The stomach was then held up by the oesophagus and completely filled with water. This volume of water was measured to the nearest centilitre.

### Transverse sections

The body was divided into 12 equal-sized, transverse sections, from the fourth gill slit (27% FL) to just anterior to the precaudal pit, with each section representing 5% of fork length (FL) (Fig 1). This was based on the procedure of Carey *et al*. (1985), and was undertaken as part of a bigger study of endothermy in this species, aimed at quantifying the amount and location of red muscle along the body. The first dorsal and pectoral fins were removed before cutting the transverse sections by hand with a knife.

**Fig. 1.**
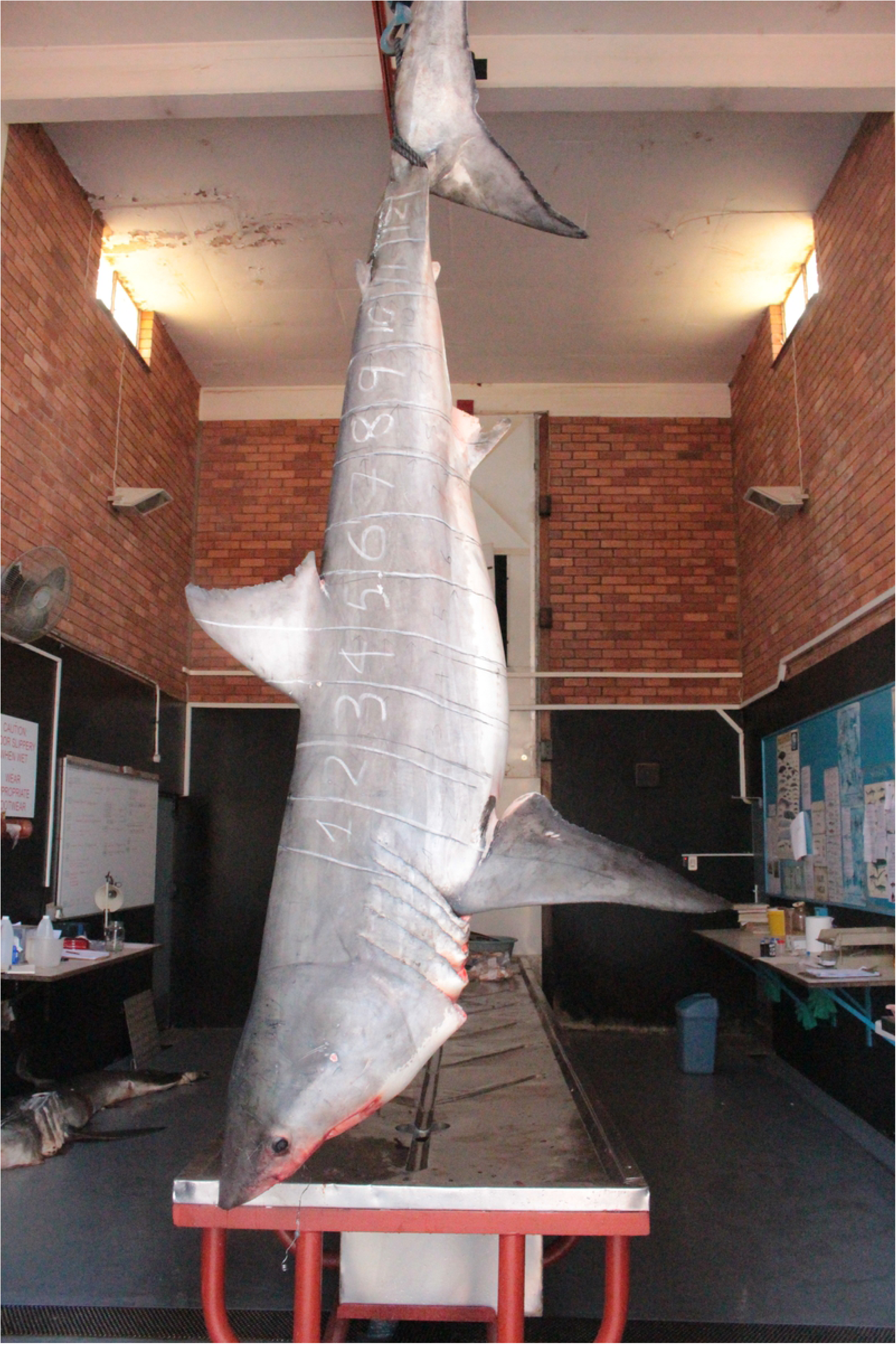
White shark (188 kg) with transverse section marks made for incisions.

The posterior ends of the transverse sections were cleaned of blood and photographed (Fig 2). The abdominal wall thickness was measured in two positions in transverse section 4 (Fig 2). A 150 kg juvenile white shark (UMH15006) was excluded due to a missing ventral abdominal wall. The thickness of the dorsolateral wall (DLW) was measured through the core of red muscles from the body cavity to the epidermis via the lateral subcutaneous vessels. This is where heat is generated through the red muscles and heat loss would be minimal compared to the thinner flanks and the belly region. As heat loss via the ventral abdomen would be greatest due to the extremely thin layer of white muscle, the ventral abdominal wall (VAW) thickness was also measured. Both measurements were made at an angle of 90° to the epidermis as a landmark (Fig 2). Student’s t-tests were used to compare VAW and DLW.

**Fig. 2.**
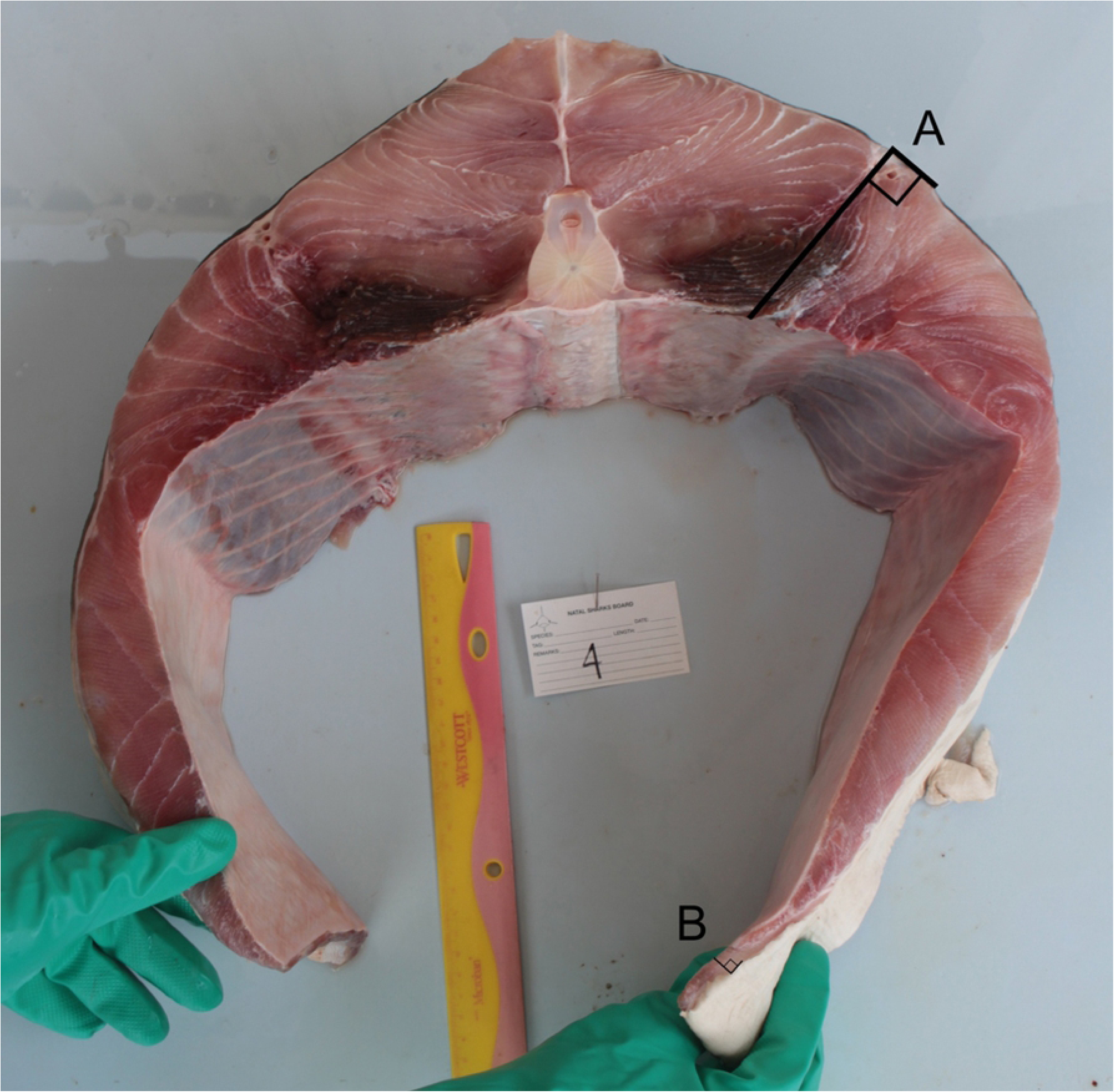
Transverse section 4 (below the first dorsal fin) from a 154 kg juvenile white shark showing body wall thickness measurements: (A) dorsolateral wall (DLW) and (B) ventral abdominal wall (VAW).

## Results

### Liver

In all white sharks examined the two liver lobes completely enveloped the entire gastrointestinal tract (GIT), with the anterio-ventral margins of the liver almost interlocking around the stomach (Fig 3). No blood vessels were evident in the outer surface of the liver. By contrast, the vessels were clearly visible on the inner surface, where they were not only closely spaced but were also flattened (Fig 4).

**Fig. 3.**
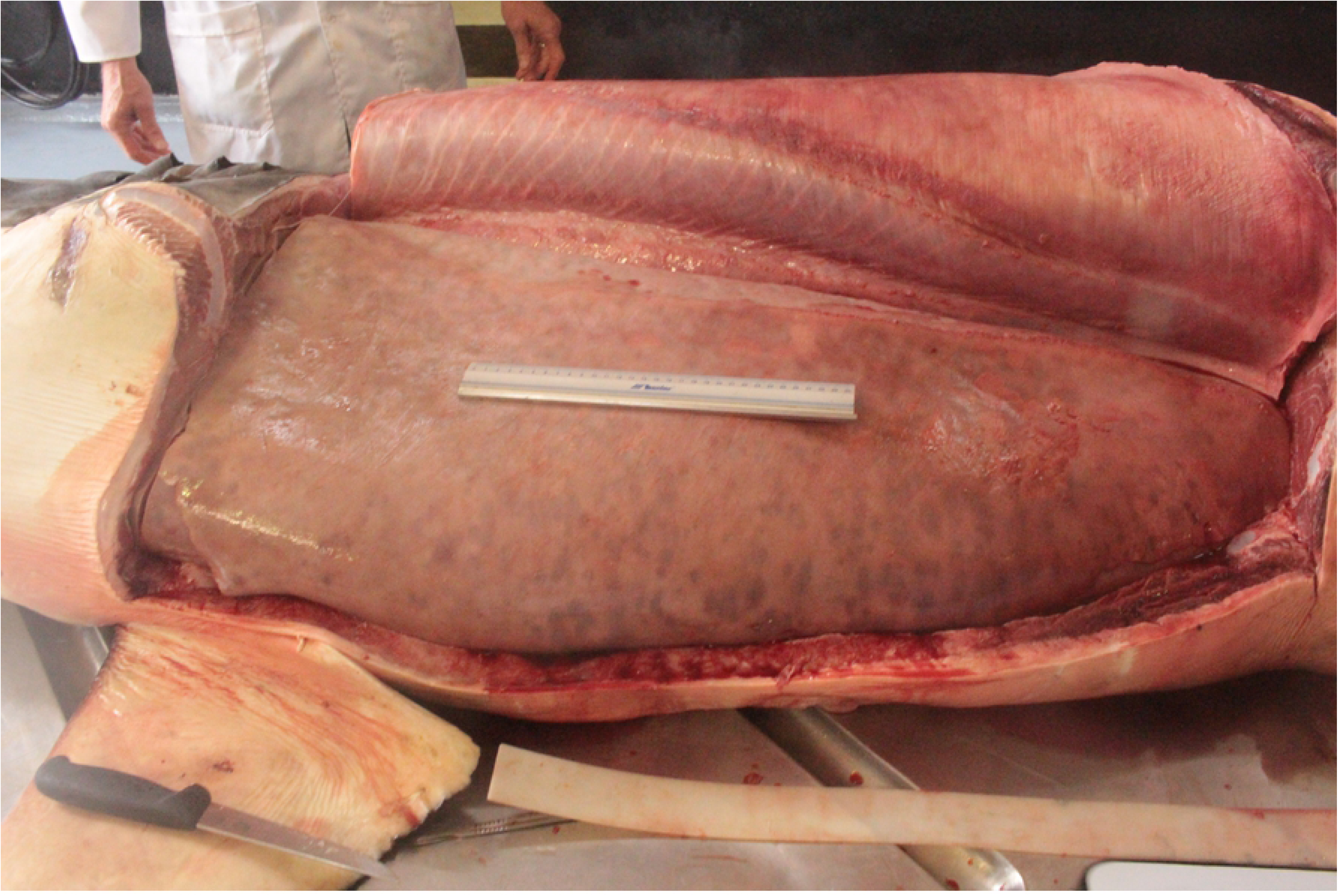
The largest white shark (296 kg). The two liver lobes completely envelope the digestive tract, keeping it away from the abdominal body wall. Note the interlocking pattern in the anterio-ventral liver margins and the absence of any vascularisation on the outer surface of the left liver lobe.

**Fig. 4.**
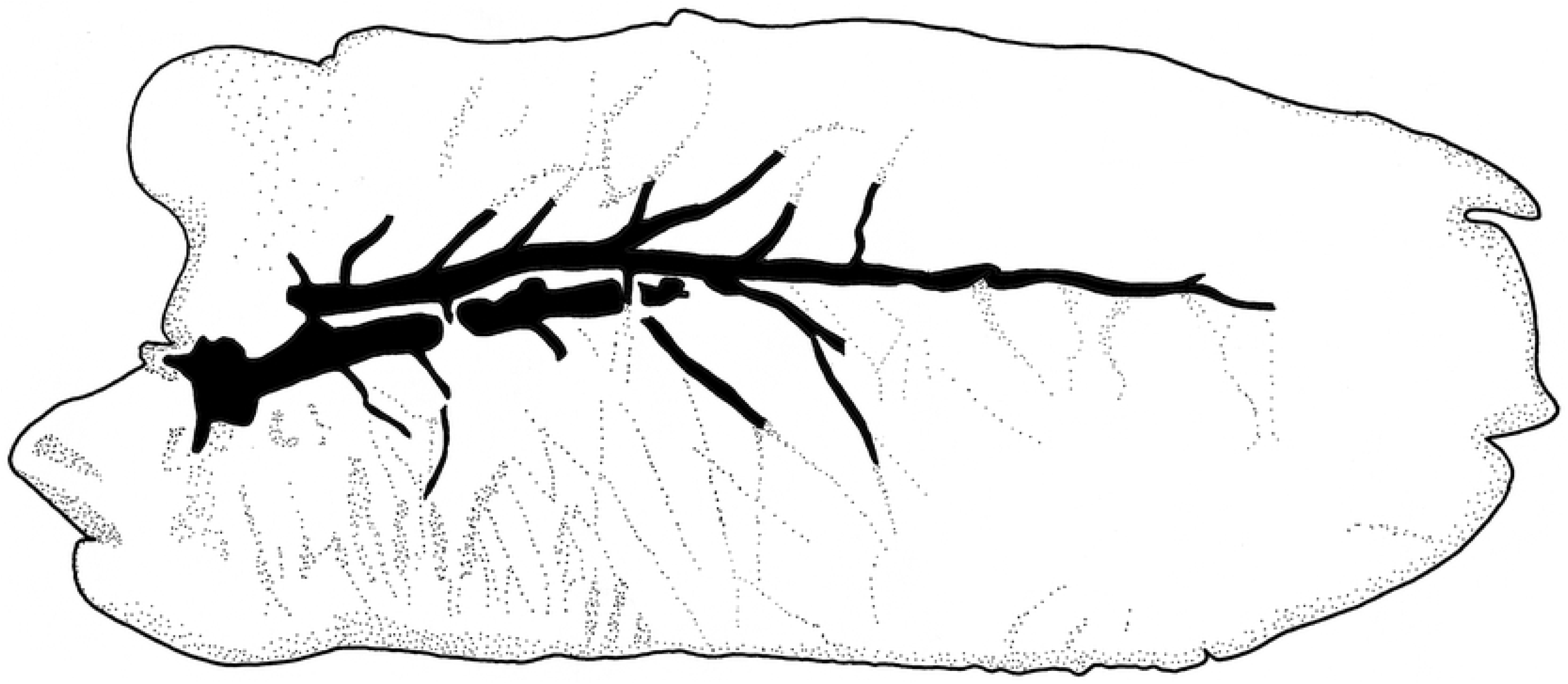
The liver lobe from a juvenile white shark (156 kg) showing pronounced vascularisation on the inside of the liver. The vessels were injected with liquid latex to make them stand out.

### Stomach volume

The stomach volume is presented in table 1, with a mean volume constituting 11.4 % of body mass (range 6.7-18.5 %, n = 11).

**Table 1:** White sharks examined in this study. KwaZulu-Natal Sharks Board identification (KZNSB-ID), total length (TL), fork length (FL), precaudal length (PCL), body mass (BM), ventral abdominal body wall for transverse section 4 (VAW), dorsolateral wall for transverse section 4 (DLW) and stomach volume (SV).

| **KZNSB-ID** | **Sex** | **TL(cm)** | **FL(cm)** | **PCL(cm)** | **BM(kg)** | **VAW(cm)** | **DLW(cm)** | **SV (l)** |
| --- | --- | --- | --- | --- | --- | --- | --- | --- |
| TRA15004 | M | 196 | 180 | 158 | 67 | 0.6 | 9.5 | 7.5 |
| LEB17007 | F | 220 | 207 | 187 | 106 | 1.6 | 9.1 | 12 |
| RB17031 | F | 224 | 209 | 185 | 105 | 1.1 | 9.8 | 18.6 |
| RB18037 | F | 239 | 215 | 195 | 130 | NA | NA | 8.7 |
| SAL18004 | M | 204 | 225 | 203 | 115 | NA | NA | 21.3 |
| UMZ15002 | M | 252 | 232 | 209 | 154 | 1.2 | 9.9 | NA |
| RB15023 | F | 255 | 234 | 210 | 126 | 0.7 | 10.4 | 12 |
| UMH15006 | F | 252 | 236 | 210 | 150 | NA | NA | NA |
| RB15017 | M | 263 | 241 | 216 | 156 | 1.1 | 10.0 | 10.9 |
| GLN17003 | M | 270 | 245 | 221 | 162 | 1.4 | 9.5 | 13 |
| MG15008 | F | 271 | 254 | 231 | 188 | 1.3 | 11.3 | 18.5 |
| LEB18004 | F | 310 | 291 | 265 | 282 | NA | NA | 31.8 |
| BAL17003 | M | 324 | 300 | 270 | 296 | 1.8 | 13.4 | 42 |

### Body wall thickness

In transverse section 4, directly below the first dorsal fin (Figure 2) and closest to the GIT, the median thickness of the VAW was 1.2 cm (range 0.6-1.4 cm, n = 9) and that of the DLW was 9.9 cm (range 9.5-13.4 cm, n = 9). As a result, the VAW below the first dorsal fin was only 12% (range 6-17.6 %, n = 9) of the DLW. There were significant positive relationships (P<0.05) between both VAW (cm) and DLW (cm) and body mass for transverse section 4 (Fig 5, Table 1).

**Fig. 5.**
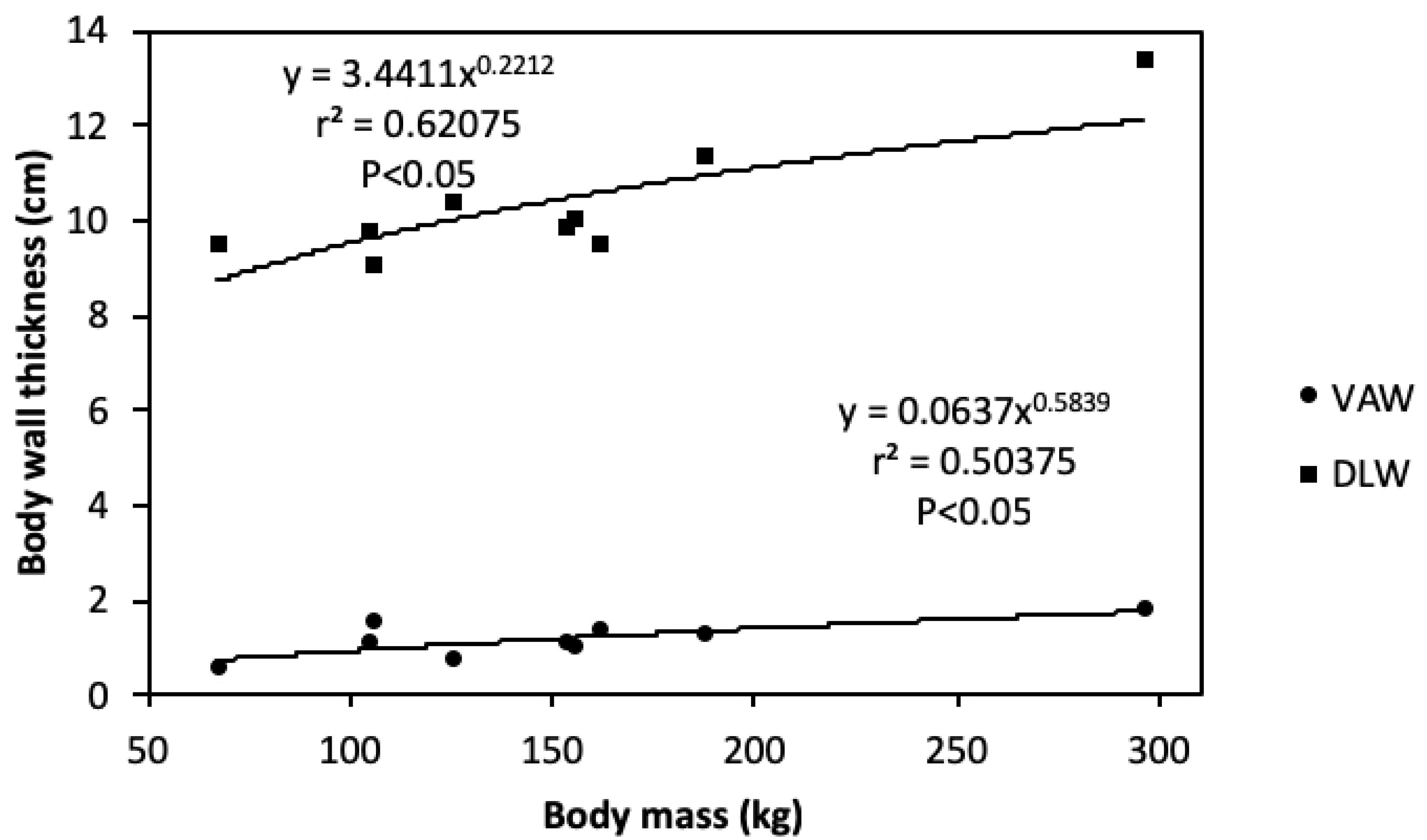
Body wall thickness below the first dorsal fin (transverse section 4) of nine white sharks showing dorsolateral wall thickness (DLW) and ventral abdominal wall thickness (VAW) in relation to body mass.

## Discussion

The current study revealed an almost total envelopment of the gastro-intestinal tract by the liver in all 13 white sharks examined, with the anterio-ventral margins of the two lobes almost interlocking. This phenomenon has not been observed in sharks of the families Carcharhinidae and Sphyrnidae (Cliff pers. comm.), but it was evident together with vascularisation on the inner liver surfaces in three shortfin mako sharks *Isurus oxyrinchus* examined during the current study. A porbeagle *Lamna nasus* examined from the North-western Atlantic Ocean showed the same pattern of vascularisation on the inner liver surfaces and envelopment of the GIT (Golet pers. comm.). This phenomenon should be investigated in other lamnid shark species i.e. longfin mako shark *Isurus paucus* and salmon shark *Lamna ditropis*. This interlocking adaptation presumably assists the liver in maintaining its enveloping role during the marked changes in stomach volume associated with the rapid ingestion of food and subsequent slow digestion. This is particularly important in a species such as the white shark which is known to ingest large prey such as dolphins, seals and the remains of baleen whales.

There are two well described sources of heat to elevate stomach temperature. The first is retention of heat generated by continuous activity of the aerobic swimming muscles and channelling it via the suprahepatic rete. This rete is well developed in all lamnids, particularly *Carcharodon* and *Lamna*, and most of the visceral blood supply passes through it (Carey *et al*. 1985). As a result, the suprahepatic rete plays a major role in warming the stomach and spiral valve (Carey *et al*. 1981). The presence of the hepatic sinus shunts between the sinus venosus and the liver allows venous blood to pass directly between these two organs but bypass the rete (Carey *et al*. 1981). We assume that warm blood is diverted to other organs while the stomach is empty, as evident in other vertebrates (Secor 2008), but when the stomach is full, blood warmed in the rete is directed to the GIT to accelerate digestion.

In addition to this source of heat, there is also the catabolic heat generated within the GIT by digestion and assimilation (Secor 2008). Our study points to a third source, that of conduction, with the warm blood from the rete flowing through prominent vessels on the inside of the liver, which is in direct contact with the GIT. The envelopment of the GIT by the liver ensures that these three sources of heat are retained, a factor which is enhanced by the presence of high concentrations of lipids similar to those found in marine mammal blubber (Davidson & Cliff 2014). These lipids not only store energy but provide insulation against heat loss. This heat loss from the GIT is potentially greatest via the ventral surface of the body, where the abdominal wall is thinnest. In the lamnids, this tissue is thinnest in white sharks and mako sharks *Isurus spp*. (Carey *et al*. 1985), which clearly does not favour heat retention. By contrast the two species of the genus *Lamna*, which frequent much colder waters, have the thickest ventral body wall of the lamnids (Carey *et al*. 1985). Heat capacity of the tissue, i.e. the thermal inertia, increases with body wall thickness (Neil *et al*. 1974).

Observations of elevated liver temperatures in *Isurus* and *Lamna* (Carey *et al*. 1985) suggest that the liver functions as a thermal insulator in all the endothermic lamnids. Furthermore, we strongly suspect that this organ envelopes the GIT in all lamnids, as it does in the three lamnid genera examined in this study. The importance of this liver envelopment and insulation increases with decreasing ambient water temperatures and abdominal body wall thickness. The hepatic sphincter muscles control blood volume and retention time of blood in the liver. When acetylcholine is administered to the hepatic sphincter muscles, contraction results and blood flow to the heart decreases (Johansen & Hanson 1967). The inference from this is that the sphincter muscles contract while the white shark is in rest-and-digest mode. As food is ingested, the rest-and-digest systemic response of the autonomous nervous system triggers a warming effect on the stomach through the hepatic shunts, directing more heated blood through the large vessels on the inside of the liver and those enveloping the GIT. There does not appear to be sufficient blood supply to the vessels of the stomach wall to greatly warm the contents, however the large, well vascularised, interlocking liver lobes could work as surrounding heat radiators to both increase the stomach temperature and conserve against heat loss through the thin abdominal wall. The liver is a very large reservoir of the blood in elasmobranchs (Johansen & Hanson 1967) and large quantities of warm blood in the white shark liver plays a major role in warming the stomach and its contents. The liver mass in the white shark may reach up to 24% of body mass (Cliff *et al*. 1989). This further demonstrates the liver is a large organ with the capacity to provide both heat and insulation to the GIT.

Heat loss is important in any endotherm which frequents cold water. In the case of white sharks which commonly ingest large prey, such as marine mammals and sharks, stomach volumes of larger individuals may exceed 100 l. A male white shark (TRA92004) from KZNSB (PCL 373 cm, BM 892 kg) had a stomach volume >135 l. The accidental ingestion of large volumes of cold sea water will rapidly lower the stomach temperature to close to that of the ambient water temperature, but it rises rapidly (within 3 minutes) to about 26°C (Goldman 1997, McCosker 1987). This rapid rise can only be facilitated by more than one mechanism, thereby resulting in enhanced rates of food digestion.

In this study we have described the almost complete envelopment of the GIT by the liver, with the two lobes interlocking along the ventral surface, and the concurrent vascularisation of the inner liver surfaces closest to the GIT. These blood vessels supply and store warm blood from the suprahepatic rete (Johansen & Hanson 1967, Carey *et al*. 1981). This provides an additional mechanism to account for the elevated stomach temperatures recorded in the wild in white sharks. The lipid-rich liver provides much needed insulation to the GIT in a region where the ventral abdominal wall is thinnest and heat loss is potentially very high. Envelopment of the GIT by a lipid-rich and well-vasculated liver facilitates the observed stomach temperatures of up to 17°C above the ambient water (Hoyos-Padilla 2016).

## Acknowledgments

We would like to thank Research and Operations staff at the KwaZulu-Natal Sharks Board, especially Kristina Naidoo, Bheki Zungu, Philip Zungu, Emmanuel Makhathini and Trish Naidu. For help with access to porbeagle liver we would like to thank Walter Golet at University of Maine and Lisa Natanson at National Oceanic and Atmospheric Administration Fisheries.

